# DNA-free CRISPR-Cas9 gene editing of tetraploid tomatoes using protoplast regeneration

**DOI:** 10.1101/2021.11.02.466947

**Authors:** Chen-Tran Hsu, Yu-Hsuan Yuan, Po-Xing Zheng, Fu-Hui Wu, Qiao-Wei Cheng, Yu-Lin Wu, Steven Lin, Jin-Jun Yue, Ying-Huey Cheng, Shu-I Lin, Ming-Che Shih, Jen Sheen, Yao-Cheng Lin, Choun-Sea Lin

**Affiliations:** Agricultural Biotechnology Research Center, Academia Sinica, Taipei, Taiwan; Biotechnology Research Center in Southern Taiwan, Academia Sinica, Tainan, Taiwan; Institute of Biochemistry, Academia Sinica, Taipei, Taiwan; Research Institute of Subtropical Forestry, Chinese Academy of Forestry, Hangzhou, China; Plant Pathology Division, Taiwan Agricultural Research Institute, Taichung, Taiwan; Department of Horticulture and Landscape Architecture, National Taiwan University, Taipei, Taiwan; Department of Molecular Biology and Centre for Computational and Integrative Biology, Massachusetts General Hospital, and Department of Genetics, Harvard Medical School, Boston, MA 02114, USA

**Keywords:** Virus resistant, DNA-free, microRNA synthesis, Ribonucleoprotein, peptide hormone

## Abstract

Wild tomatoes are important genomic resources for tomato research and breeding. Development of a foreign DNA-free CRISPR-Cas delivery system has potential to mitigate public concern about genetically modified organisms. Here, we established a DNA-free protoplast regeneration and CRISPR-Cas9 genome editing system for *Solanum peruvianum*, an important resource for tomato introgression breeding. We generated mutants for genes involved in small interfering RNAs (siRNA) biogenesis, *RNA-DEPENDENT RNA POLYMERASE 6* (*SpRDR6*) and *SUPPRESSOR OF GENE SILENCING 3* (*SpSGS3*); pathogen-related peptide precursors, *PATHOGENESIS-RELATED PROTEIN-1* (*SpPR-1*) and *PROSYSTEMIN* (*SpProsys*); and fungal resistance (*MILDEW RESISTANT LOCUS O, SpMlo1*) using diploid or tetraploid protoplasts derived from *in vitro*-grown shoots. The ploidy level of these regenerants was not affected by PEG-calcium-mediated transfection, CRISPR reagents, or the target genes. By karyotyping and whole genome sequencing analysis, we confirmed that CRISPR-Cas9 editing did not introduce chromosomal changes or unintended genome editing sites. All mutated genes in both diploid and tetraploid regenerants were heritable in the next generation. *spsgs3* null T_0_ regenerants and *sprdr6* null T_1_ progeny had wiry, sterile phenotypes in both diploid and tetraploid lines. The sterility of the *spsgs3* null mutant was partially rescued, and fruits were obtained by grafting to wild-type stock and pollination with wild-type pollen. The resulting seeds contained the mutated alleles. Tomato yellow leaf curl virus proliferated at higher levels in *spsgs3* and *sprdr6* mutants than in the wild type. Therefore, this protoplast regeneration technique should greatly facilitate tomato polyploidization and enable the use of CRISPR-Cas for *S. peruvianum* domestication and tomato breeding.

**One-sentence summary:** DNA-free CRISPR-Cas9 genome editing in wild tomatoes creates stable and inheritable diploid and tetraploid regenerants.

## Introduction

Tomato is an important vegetable crop, representing the sixth most economically important crop worldwide (http://www.fao.org/faostat/en/#data/QV). Wild tomato species are resistant to diverse biotic and abiotic stresses, and are often used for tomato introgression breeding. De novo domestication of wild tomato was recently achieved within a short period by gene editing using clustered regularly interspaced short palindromic repeats (CRISPR) and CRISPR-associated protein (Cas) (Li et al., 2018; Zsogon et al., 2018). Thus, CRISPR-Cas mutagenesis of wild tomato represents a new strategy for tomato breeding and basic research.

Genome multiplication is a frequent occurrence during crop domestication. Many of the most economically important crops are polyploid, including potato, wheat, and cotton. Polyploidy conveys advantages in terms of genomic buffering, viability, and environmental robustness (Van de Peer et al., 2021). Triploids can also be used as seedless crops, such as watermelon and bananas. Thus, CRISPR-Cas-edited tetraploid versions of crop species and their relatives represent important materials for crop breeding in the face of rapid climate change caused by global warming, among other challenges, as was recently demonstrated for tetraploid wild rice (*Oryza alta*) (Yu et al., 2021). Therefore, it is important to establish a gene editing platform for polyploid crops and related species.

The CRISPR-Cas system uses *Agrobacterium*-mediated stable transformation to deliver DNA encoding Cas protein and single guide RNA (sgRNA) into the nuclei of tomato cells. As an alternative approach, CRISPR ribonucleoprotein (RNP) or plasmids harboring the Cas and sgRNA sequences can be introduced directly into protoplasts using transient transfection, allowing recombinant DNA-free plants to be regenerated to circumvent concerns about genetically modified organisms (GMOs) (Woo et al., 2015; Andersson et al., 2018; Lin et al., 2018; Hsu et al., 2019; De Bruyn et al., 2020; Hsu et al., 2021; Hsu et al., 2021; Yu et al., 2021). This protocol is important for use with hybrids or plants with a long juvenile period and for vegetative propagation because the transgenes from stable transformation (selection markers and CRISPR reagent genes) cannot be removed from these crops by crossing. Also, the progeny will be different from their heterozygous parental lines due to segregation. The gene editing efficiency and specificity could be validated by targeted sequencing (Woo et al., 2015; Nekrasov et al., 2017) or whole genome sequencing (WGS) (Fossi et al., 2019; Hsu et al., 2021). Nevertheless, previous analysis paid little attention to the overall chromosomal changes, especially in polyploid regenerants (Fossi et al., 2019).

The protoplast regeneration gene editing system has two other major advantages: (1) Gene-edited transformants derived from tissue-culture-based *Agrobacterium*-mediated transformation are often chimeric, especially in dicotyledons (Shimatani et al., 2017). If the transformant is an edited/wild-type (WT) chimera and the edited allele occurs only in somatic cells (and not germ cells), edited alleles cannot be passed on to the next generation (Zheng et al., 2020). In protoplast regeneration, there is a low incidence of chimerism, and all mutated alleles detected in the T_0_ generation can be transmitted to the next generation (Lin et al., 2018; Hsu et al., 2019; Hsu et al., 2021; Hsu et al., 2021). (2) The protoplast regeneration system can be used to introduce many CRISPR reagents and donor DNAs into plants for targeted insertion at the same time without the limitation of vector size (Hsu et al., 2019; Hsu et al., 2021). In addition, the second transfer step can be performed directly to obtain homozygous alleles in polyploids without self-fertilization which is very useful for hybrid, long juvenile period, and sterile plants (Hsu et al., 2019). However, the main bottleneck of this strategy is the difficulty of performing protoplast regeneration.

Here, we established a diploid/allotetraploid protoplast regeneration protocol for *S. peruvianum*, an important stress-resistant wild tomato, for use with CRISPR-Cas-mediated genome editing. We targeted several genes for editing, including *RNA-DEPENDENT RNA POLYMERASE6* (*SpRDR6*) and *SUPPRESSOR OF GENE SILENCING3* (*SpSGS3*), two key genes in the plant RNA silencing pathway (Mourrain et al., 2000) that mediate defense against tomato yellow leaf curl virus (TYLCV) (Verlaan et al., 2013); *PATHOGENESIS-RELATED PROTEIN-1* (*SpPR-1*) encoding the cysteine-rich secretory proteins antigen 5 and pathogenesis-related 1 protein (CAP)-derived peptide 1 (CAPE1) precursor (Chen et al., 2014) and *PROSYSTEMIN* (*SpProsys*), two pathogen-resistance peptide precursors; and *MILDEW RESISTANT LOCUS O* (*SpMlo1*) (Nekrasov et al., 2017). Targeting of these genes, which was performed using two types of CRISPR reagents, plasmids and RNPs, yielded diploid and tetraploid transgene-free lines. Stable genome structures of ten plants, including one explant derived from stem cutting, three diploid regenerants and six tetraploid of *SpProsys* or *SpMlo1* RNP transfection regenerants were confirmed by WGS.

## Results

### Protoplast regeneration in *S. peruvianum*

To obtain a high proportion of tetraploid protoplasts, we analyzed the genome sizes of different explants (leaves and stems) using flow cytometry to determine the proportion of tetraploid cells. In leaves, the ratio of diploid to tetraploid nuclei was 5:1 (Figure 1a), and in stems, the ratio was 1:1 (Figure 1b). The same ratio was detected in protoplasts derived from stems (Figure 1c). Therefore, since stems had a higher proportion of tetraploid cells, we used them in subsequent studies to increase the proportion of tetraploid regenerated plants.

**Figure 1.**
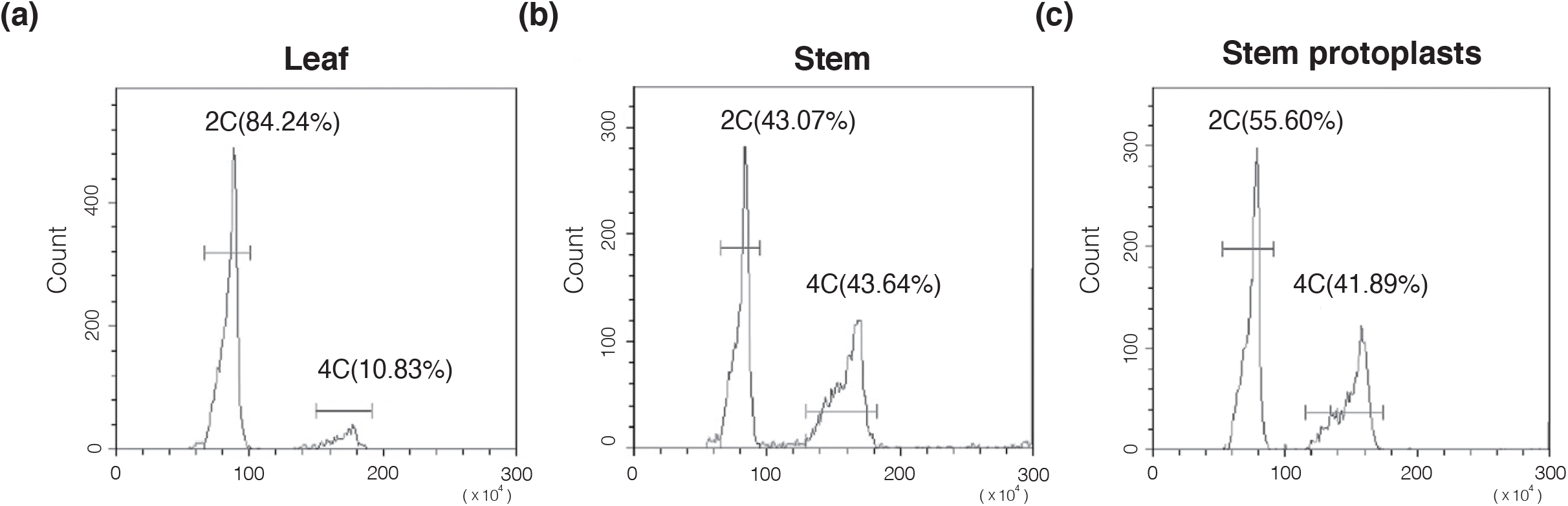
Flow cytometric analysis of the nuclear DNA contents of *S. peruvianum* tissues. The genome sizes of **(a)** leaves, **(b)** stems **(c)**, and protoplasts derived from stems. X: fluorescence density; Y: count. Chicken erythrocyte nuclei (CEN: 2.5 Gb) were used as the calibration standard. The bar indicates the area used for counting nuclei. 2C: diploid; 4C: tetraploid. The number in brackets after the ploidy is the percentage of each different ploidy level versus the total counts.

Using a method previously published for *Nicotiana tabacum* (Lin et al., 2018), we successfully isolated *S. peruvianum* protoplasts from *in vitro-*grown shoots. We incubated the purified protoplasts in liquid medium consisting of half-strength Murashige and Skoog medium (1/2 MS), 0.4 M mannitol, 3% sucrose, 1 mg/L naphthaleneacetic acid (NAA), and 0.3 mg/L kinetin, pH 5.7, for 1 month in the dark, leading to the formation of fine, sand-like calli (Figure 2a). Next, we subcultured these calli in liquid medium containing 1/2 MS, 0.4 M mannitol, 3% sucrose, 2 mg/L kinetin, and 0.3 mg/L Indole-3-acetic acid (IAA), pH 5.7, in the light (Figure 2b). After one month, these white calli turned green and were transferred to solid medium (1/2 MS, 0.2 M mannitol, 3% sucrose, and 2 mg/L kinetin; Figure 2c). We transferred the calli to fresh medium every month to induce the formation of small shoots (Figure 2d), which were incubated in medium without plant growth regulators until adventitious roots formed at the bottoms of the shoots (Figure 2e). Finally, we transferred the rooted plants to pots (Figure 2f) and grew them in the greenhouse (Figure 2g). The regenerated plants flowered (Figure 2h), fruited (Figure 2i), and produced seeds.

**Figure 2.**
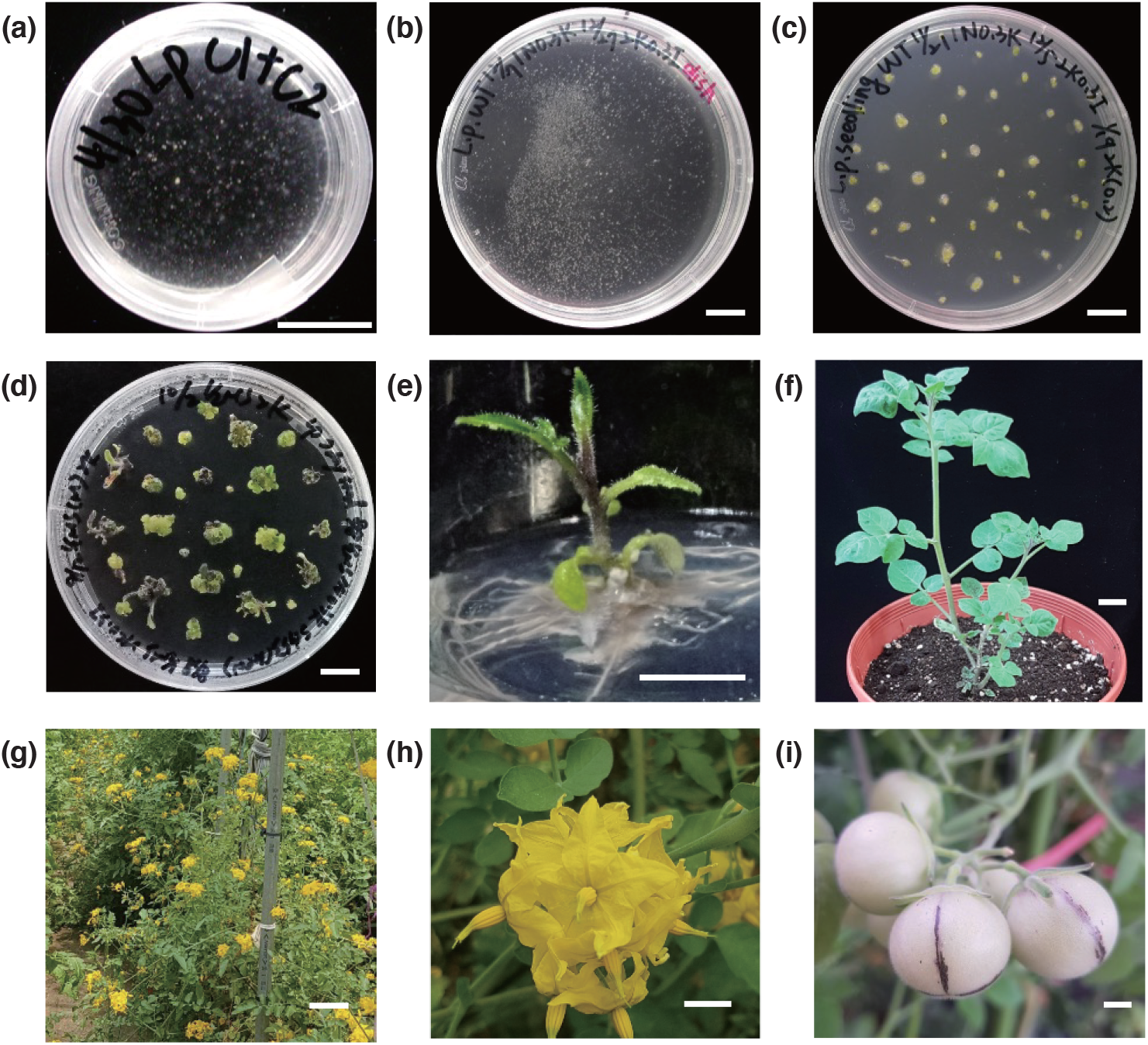
Regeneration of *S. peruvianum* protoplasts. Protoplasts incubated in 1/2 Murashige and Skoog (MS) medium supplemented with 3% sucrose, 0.4 M mannitol, 1 mg/L naphthaleneacetic acid (NAA), and 0.3 mg/L kinetin, pH 5.7 liquid medium for 1 month. **(b)** Calli subcultured in 1/2 MS medium supplemented with 3% sucrose, 0.4 M mannitol, 2 mg/L kinetin, and 0.3 mg/L Indole-3-acetic acid (IAA), pH 5.7 liquid medium in the light. **(c)** Calli subcultured in 1/2 MS medium supplemented with 3% sucrose, 0.4 M mannitol, 2 mg/L kinetin, pH 5.7, solid medium. **(d)** Shoot bud formation after two subcultures in 2 mg/L kinetin solid medium. **(e)** Adventitious root formation in plant growth regulator-free 1/2 MS solid medium supplemented with 3% sucrose. **(f)** Regenerated plants after 1 month of growth in a pot. **(g)** Regenerated plants grown in the field. **(h)** Flowers of a regenerated plant. **(i)** Fruits of a regenerated plant. Throughout, bars = 1 cm.

### Optimized protoplast regeneration protocol

Compared to tobacco (Lin et al., 2018), *S. peruvianum* protoplasts take longer to regenerate. According to our observations, the most important steps in the tomato regeneration process are those in liquid culture: callus induction in the dark (the 1^st^ step) and callus proliferation in the light (the 2^nd^ step). Therefore, we tested several modifications to the composition of the culture medium to shorten the regeneration time. The results indicated that zeatin and 6-Benzylaminopurine (BA) are the best hormonal treatments for the two liquid culture steps (Figure S1), and zeatin is the best cytokinin for the 3^rd^ subculture step in solid medium (Figure S2).

### CRISPR-Cas9-targeted mutagenesis in S. *peruvianum*

We used this protoplast regeneration system to establish a method for CRISPR-Cas9-targeted gene mutagenesis of *S. peruvianum*. First, we used plasmids as CRISPR-Cas9 reagents for targeting mutagenesis of three important disease-resistance-related genes: *SpSGS3, SpRDR6*, and *SpPR-1*.

In the *SpSGS3* experiment, we chose four target sites (Table 1), and the total efficiency of mutagenesis was 8.3%. Based on sequencing results, mutations occurred in all three target sites except GTAACAATGCTGGATCAGGC. Among these, GCGCAATTGAATGGTTTACA was targeted the most effectively, and mutations at this position were observed in all mutants (Table S1). *spsgs3*#6, #11, and #13 are null mutants and *spsgs3*#6 contains four mutated alleles. *SpSGS3*#7 also contains three mutated alleles and one non-mutated WT allele. A 68-bp insertion from the vector was detected in *spsgs3*#11.

**Table 1.**
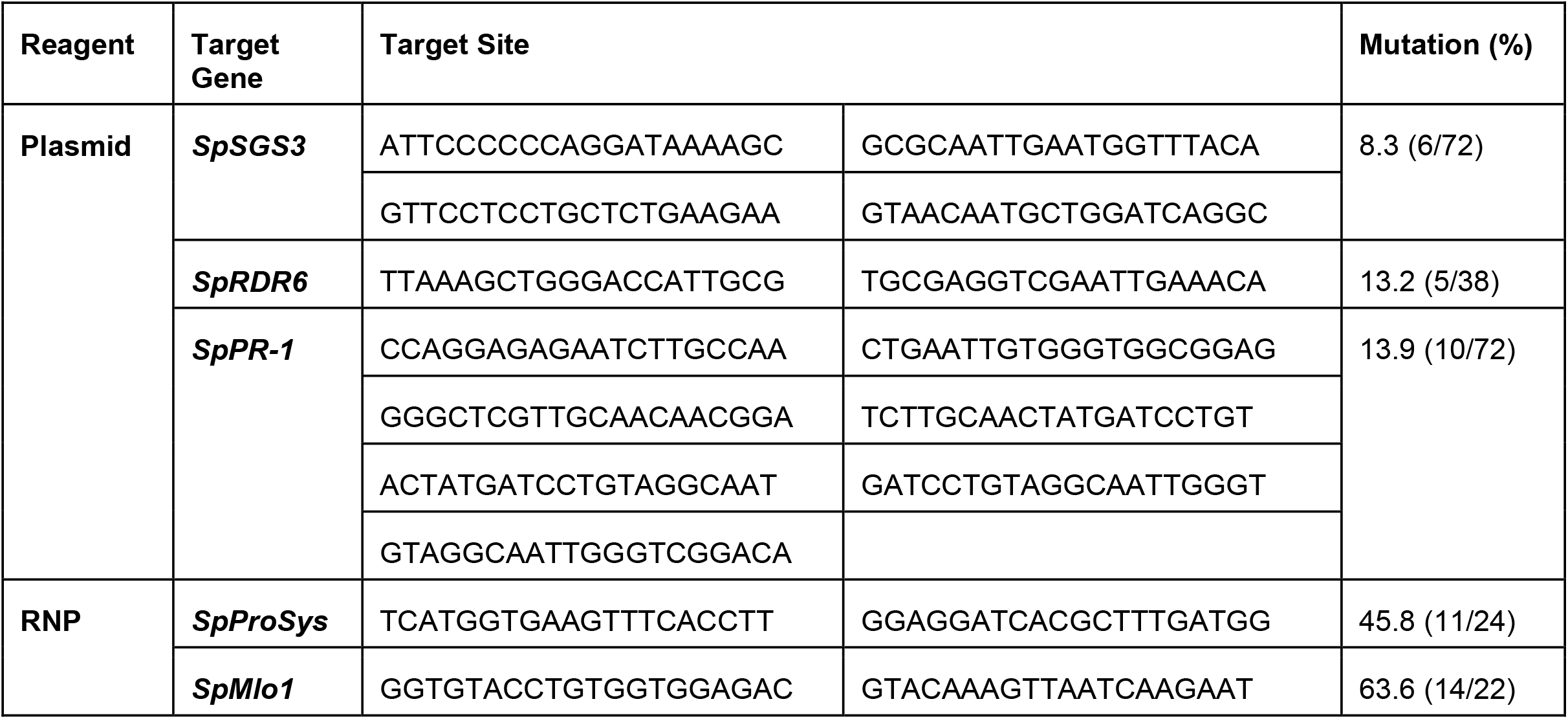
CRISPR-Cas9 target sites and mutagenesis efficiencies.

In the *SpRDR6* experiment, we selected two target sites (Table 1). Based on the sequencing results, both target sites could be mutated by CRISPR-Cas9, with a total mutation efficiency of 13.2%. TTAAAGCTGGGACCATTGCG gave the best results, as all five mutant plants contained mutations at this target site. The mutation TGCGAGGTCGAATTGAAACA was only identified in *SpRDR6*#38 (Table S2). All regenerated mutants were heterozygous, and *SpRDR6*#38 had two mutated alleles and at least one WT allele.

In the *SpPR-1* experiment, seven target sites were selected and used to construct two vectors. These two constructs, harboring sgRNAs targeting seven target sites, were co-transfected into protoplasts (Table 1). Among the 10 regenerated mutants, 4 contained fragment deletions, indicating that at least two cleavages had occurred. Except for TGTCCGATCCAGTTGCCTAC and CTATGATCCTGTAGGCAAC there were no mutations in the target sites; the five other sgRNAs caused mutations at the expected positions. The mature CAPE1 peptide is derived from the C-terminal end of tomato *PR-1b. sppr-1*#28, #31, and #52 were mutated only in the target sites located in CAPE1, all at ATCCTGTAGGCAACTGGAT, resulting in a 5-bp deletion. All *SpPR-1* mutants were null mutants except for *SpPR-1*#72 (Table S3).

In the experiments with *N. tabacum* (Lin et al., 2018) and *SpSGS3* (Table S1), the use of plasmid CRISPR reagent may still result in foreign DNA insertions. Therefore, RNP is used as a CRISPR reagent to achieve DNA-free gene editing. Here, we delivered two RNPs that target sites located in *SpProsys* to protoplasts and regenerated the transfected protoplasts into plants. Upon sequencing of the 24 regenerated *SpProsys* plants, 11 showed target mutagenesis (45.8%, Table 1). Prosystemin is a precursor of systemin, which is processed by phytaspase (Beloshistov et al., 2018). The target site GGAGGATCACGCTTTGATGG is at the C terminus of SpProsys, which is the position of systemin, and the mutations in lines #5, #16, and #19 occurred only at this site (Table S4). Using two published *SlMlo1* target sites (Nekrasov et al., 2017), we synthesized RNPs targeted to these sites *in vitro* and simultaneously delivered them into protoplasts. Of the regenerated calli and plants, 63.6% showed targeted mutagenesis (Table 1, Table S5).

### Analysis of the genome sizes, phenotypes, and progeny of diploids and tetraploids

A higher proportion of tetraploid cells was observed in protoplasts derived from diploid stems compared to leaf tissue (Figure 1). In addition, during target gene genotyping, we observed that some mutants contained more than three alleles. For example, *SpRDR6*#38 contained three alleles (+1 bp, –7 bp, and WT, Table S2), and its genome size was 4.40 ± 0.03 pg. Therefore, targeted mutant plants of tetraploids can be obtained using this method. We performed karyotype analysis of these regenerated plants (T_0_, sterile mutants) or their offspring (T_1_) to confirm the chromosome numbers (Figure 3). Except for a *SpPR-1* tetraploid without targeting regenerant, we obtained diploid and tetraploid regenerated plants with or without targeting mutations derived from plasmid CRISR-Cas9 reagent-transfected protoplasts (Figure 3, Table S6). Similar results were obtained for *SpProsys* RNP transfection (Figure S3, Table S7). The ploidy of the plants that were regenerated from transfected protoplasts is provided in Table S6 and S7. These results indicate that most tetraploid plants were derived from tetraploid protoplasts from the explants rather than by protoplast fusion caused by the presence of PEG-Ca^+^ in the transfection medium.

**Figure 3.**
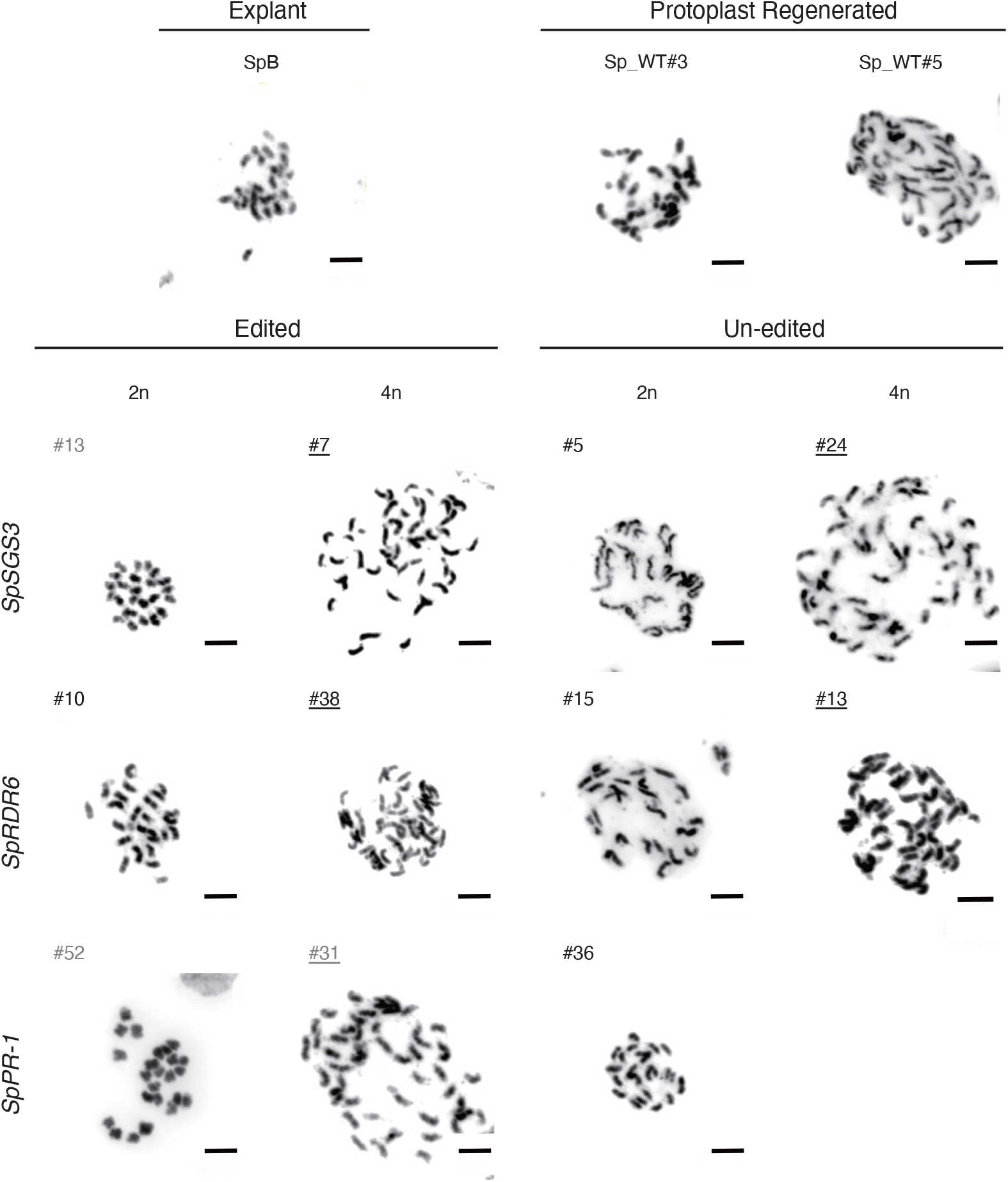
Karyotypes of *S. peruvianum* plants regenerated from protoplasts. Gray font: null mutant. Black font: heterozygous or wild-type. Underline: 4n. Bars = 5 μm.

In regenerated plants derived from *SpSGS3* transfection, the tetraploids had a reduced seed set (Figure S4a). The seeds of tetraploids were larger than those of diploids; this phenomenon was also observed in tetraploid regenerated plants derived from transfection with other CRISPR reagents (Figure S4b). The tetraploid plants grew more slowly than the diploid plants (Figure S4c). The leaf edges of tetraploid plants were more rounded than those of the diploid plants (Figure S4c).

We subjected the offspring of *SpSGS3*#7, and #10 (Figure S5); *SpRDR6*#6, #33, and #38 (Figure S6); and *sppr-1*#52 and #61 (Figure S7) to target gene sequencing. Except for *sppr-1*#52, which contained one mutant locus not present in the parent, all other offspring had the same mutated locus as the parent. These results demonstrate that these mutated loci can be transmitted to the next generation in diploids and tetraploids.

### Stable genome structures in diploids and tetraploids

To further confirm the stability of genome structure in regenerants, we performed whole genome sequencing of ten samples, including one diploid plant propagated by stem cutting (SpB), three diploids and six tetraploids derived from *SpProsys* or *SpMlo1* RNP transfection (Table S7). Taking into account the different genome sizes between diploid and tetraploid plants, each sample was sequenced to the anticipated 30x genome coverage. That is, 141-171 million pair-end reads were sequenced for diploid plants and at least 252-373 million pair-end reads were sequenced for tetraploid plants (Table S7).

Multiple analysis strategies were used to study the genome structures. Despite the low mapping rate of both diploid and tetraploid samples at some chromosome locations, sequencing coverage analysis did not show inconsistent coverage changes between samples (Figure S8). Deletion of large chromosomal segments, which were commonly seen in aneuploid cells (Musacchio and Salmon, 2007) cause allelic imbalance. By calculating heterozygous allele frequency of sequenced plants, we did not identify abnormal allele frequency variations or loss of heterozygosity (Figure 4a). A Bayesian approach to determine copy number variations along chromosomes and compared between uneven sequencing depth of samples did not identify abnormal copy number changes in sequenced plants (Figure 4b). Taking these findings together, we concluded that there is no abnormal chromosomal gain or loss in diploid and tetraploid plants. Neither the protoplast regeneration process nor the CRISPR reagents caused detectible chromosomal changes.

**Figure 4.**
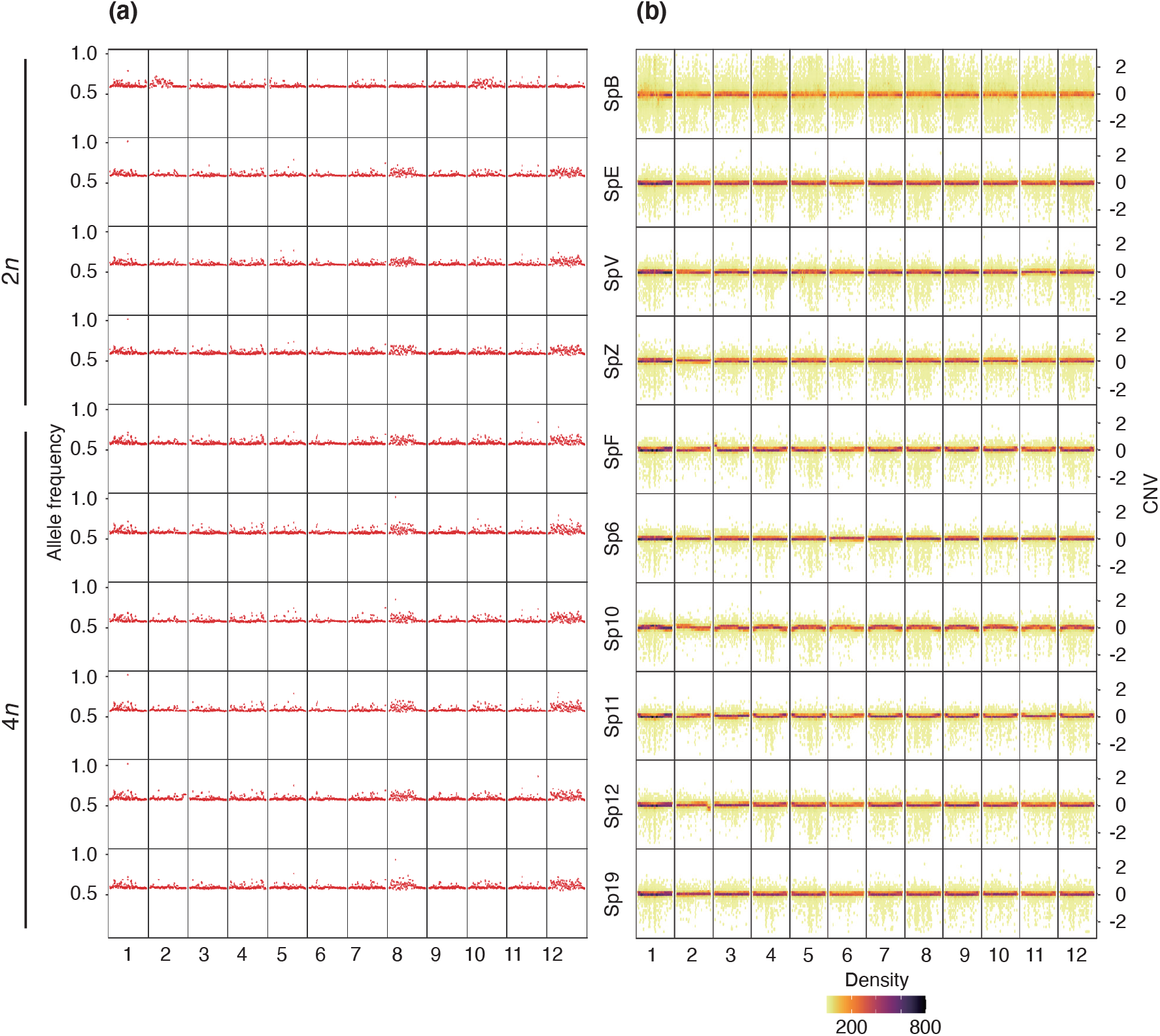
Stable genome structures in plants regenerated from stem cutting and protoplasts. **(a)** Heterozygous allele frequency of WGS samples. The heterozygous allele frequency was attained by dividing the read depth of the heterozygous allele (labeled as 0/1 by GLnexus) by the total read depth of the variant. Heterozygous frequency is plotted using 10-kb chromosome window size on the X axis. A value of heterozygous allele frequency 0.5 indicates the frequency of the heterozygous genotype (0/1) from the DeepVariant is 0.5, regardless the ploidy level. **(b)** Copy number variation (CNV) of WGS samples. CNV was predicted as 3kb fragment size with minimum 10 fragments. Predicted CNV is plotted using 30 bins per chromosome on the X axis. Dot colors indicate the CNV density per bin. A value of zero on the Y axis indicates no copy number change was detected. Values above zero indicate copy number gain and below zero indicate copy number loss.

### *spsgs3* and *sprdr6* diploid and tetraploid null mutants show wiry phenotypes

The regenerated plants containing a WT allele(s) produced flowers and fruits (Figure 2) with morphology and development similar to those of WT plants in the greenhouse. Biallelic *spsgs3* mutants (carrying two distinct genome-edited alleles: *spsgs3*#11, Figure 5; *spsgs3-6* and *spsgs3-13*; Figure S9a) had a wiry leaf phenotype and abnormal flowers, which is similar to the previously reported *sgs3* domesticated tomato mutants (Yifhar et al., 2012).

**Figure 5.**
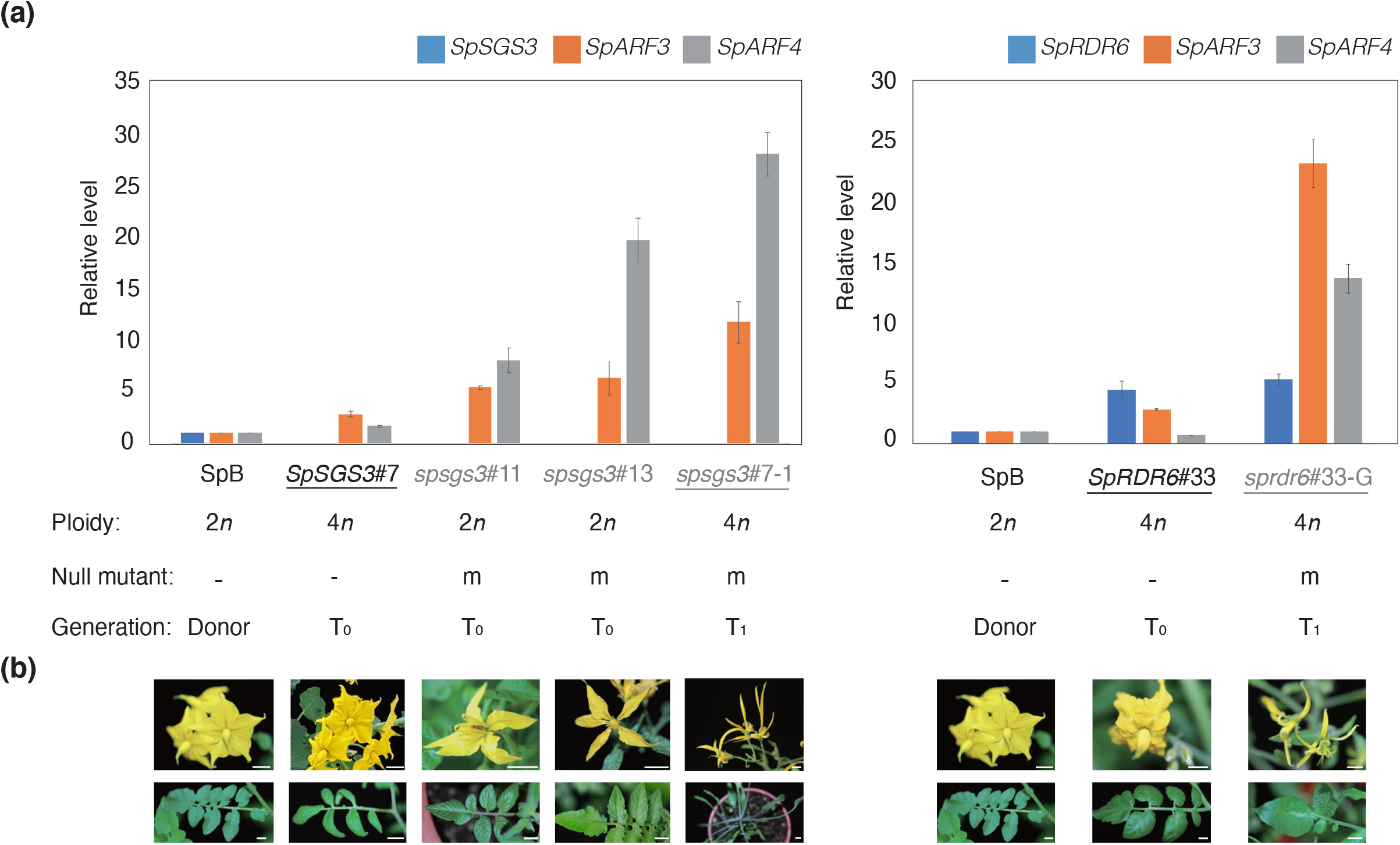
Gene expression and phenotypic profiles of *S. peruvianum sgs3* and *rdr6* mutants. **(a)** RT-qPCR analysis of auxin response regulator genes (*SpSGS3, SpARF3, SpARF4* and *SpRDR6*) in the wild type and protoplast-derived regenerants. T_0_: regenerated plants derived from protoplasts. T_1_: seedlings derived from T_0_ plants. **(b)** Phenotypes of *spsgs3* and *sprdr6* mutants. Bars = 1 cm.

Among the six progeny of *SpSGS3*#7, two progeny harbored the mutated alleles only (Figure S5); these plants also showed a wiry phenotype (*spsgs3*#7-2; Figure S9b). A similar phenomenon was also observed in the *SpRDR6* regenerants. Although all *SpRDR6* T_0_ plants were heterozygous and contained WT *SpRDR6* alleles in their genomes, no wiry phenotypes were observed. The *SpRDR6*#33 and *SpRDR6*#38 offspring had wiry phenotypes (*sprdr6*#33-G, Figure 5b; *sprdr6*#38-16, Figure S9b). The pollen of both null T_0_ and T_1_ mutated plants, including *SpSGS3* and *SpRDR6* mutants, was abnormal (Figure S9c) and failed to produce seeds.

Because *AUXIN RESPONSE FACTOR3* (*ARF3*) and *ARF4* are the target genes of trans-acting secondary siRNA3 (*TAS3*), whose biogenesis requires RDR6 (Marin et al., 2010), we investigated the transcript levels of these genes in WT, *spsgs3* and *sprdr6* plants (Figure 5a). The *spsgs3* null mutants (T_0_: *spsgs3*#11 and *spsgs3*#13; T_1_: *spsgs3*#7-1) lacked *SpSGS3* expression. In contrast to the WT, the transcript levels of *SpARF3* and *SpARF4* was increased in the *spsgs* mutants, not only for null diploid mutants *spsgs3*#11 and *spsgs3*#13 but also for tetraploid mutant *spsgs3*#7-1 (Figure 5a). Similarly, the transcript levels of *SpARF3* and *SpARF4* were also increased in the *SpRDR6* T_1_ mutant *sprdr6*#33-G (Figure 5a).

### Tomato yellow leaf curl virus proliferation

We evaluated the infectivity of TYLCV in the mutants by *in vitro* inoculation (Al Abdallat et al., 2010). After 8 weeks *in vitro* inoculation, plant growth was severely retarded (Figure S10) and leaf morphology changed in the T_1_ diploid *spsgs3*#11 (Figure 6a) and the T_2_ tetraploid *sprdr6*#38-6 (Figure 6b). Compared to the WT, all of the null mutants (*spsgs3*/*sprdr6* and diploid/tetraploid) showed higher levels of TYLCV accumulation (Figure 6a, b, Figure S10).

**Figure 6.**
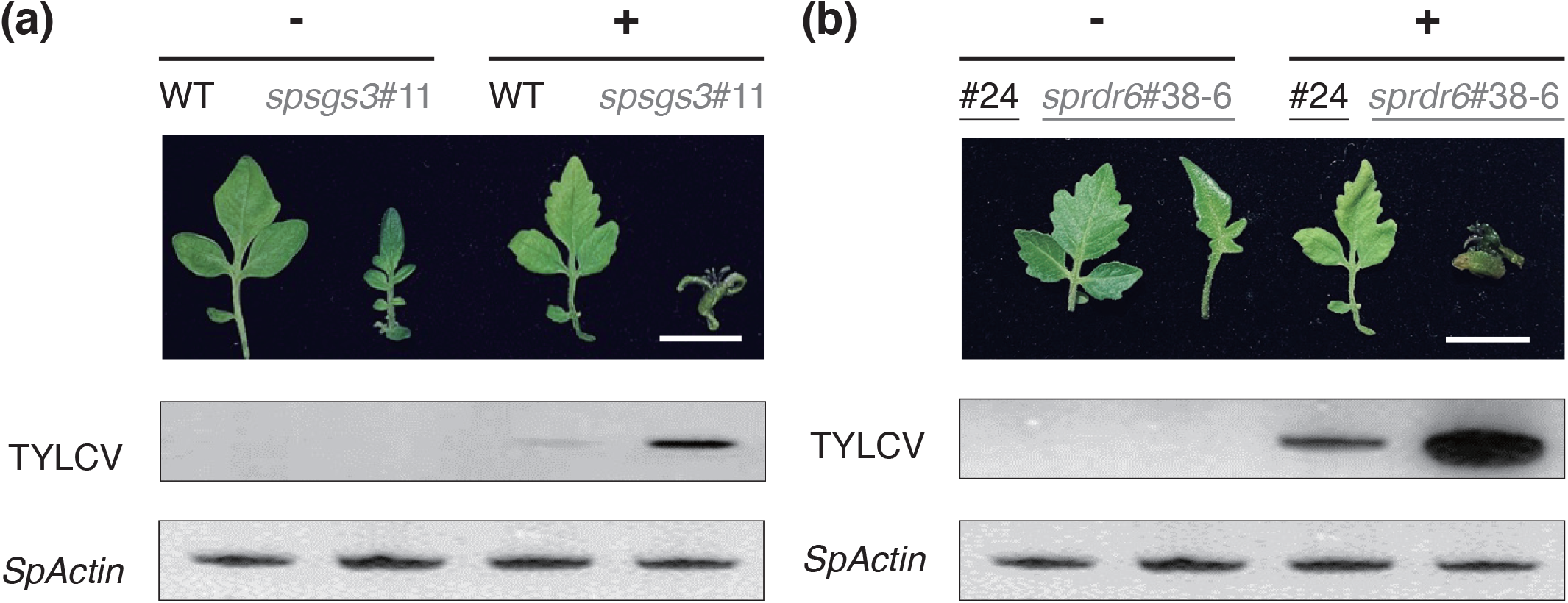
Symptoms and TYLCV proliferation on *in vitro*-cultured *S. peruvianum* plants inoculated with the infectious TYLCV clone. **(a)** Diploid wild type and *spsgs3#11* mutant. **(b)** Tetraploid regenerated plant (#24) and *sprdr6#38-6* mutant. Gray: null mutant. Black: Un-edited tetraploid regenerated plant (#24) or the wild type. Underline: 4n. Bars = 1 cm.

### Grafting rescued the fertility of the *sgs3#*11 null mutant

We used WT pollen for hybridization, which failed to pollinate the fruits of the *spsgs3* and *sprdr6* null mutants. Based on these results, these mutants could not produce the substrate(s) needed for the development of male or female reproductive organs. However, using grafting, the substrate(s) produced in WT stock was successfully transported to the *spsgs3*#11 scion (Figure 7a). Although there were no significant differences in leaf (Figure 7b) or flower morphology (Figure 7c), *spsgs3*#11 failed to produce viable pollen (Figure S9c) and the pollen viability of *spsgs3*#11 increased to 20% by grafting to the WT stock. Grafted *spsgs3*#11 produced fruits (Figure 7d), but non-grafted *spsgs3*#11 did not. The fruits from *spsgs3*#11 scions were smaller (Figure 7e) and contained fewer seeds than the WT (Figure 7f). Genotyping indicated that all of the progeny harbored *spsgs3*#11 mutated alleles (Figure 7g).

**Figure 7.**
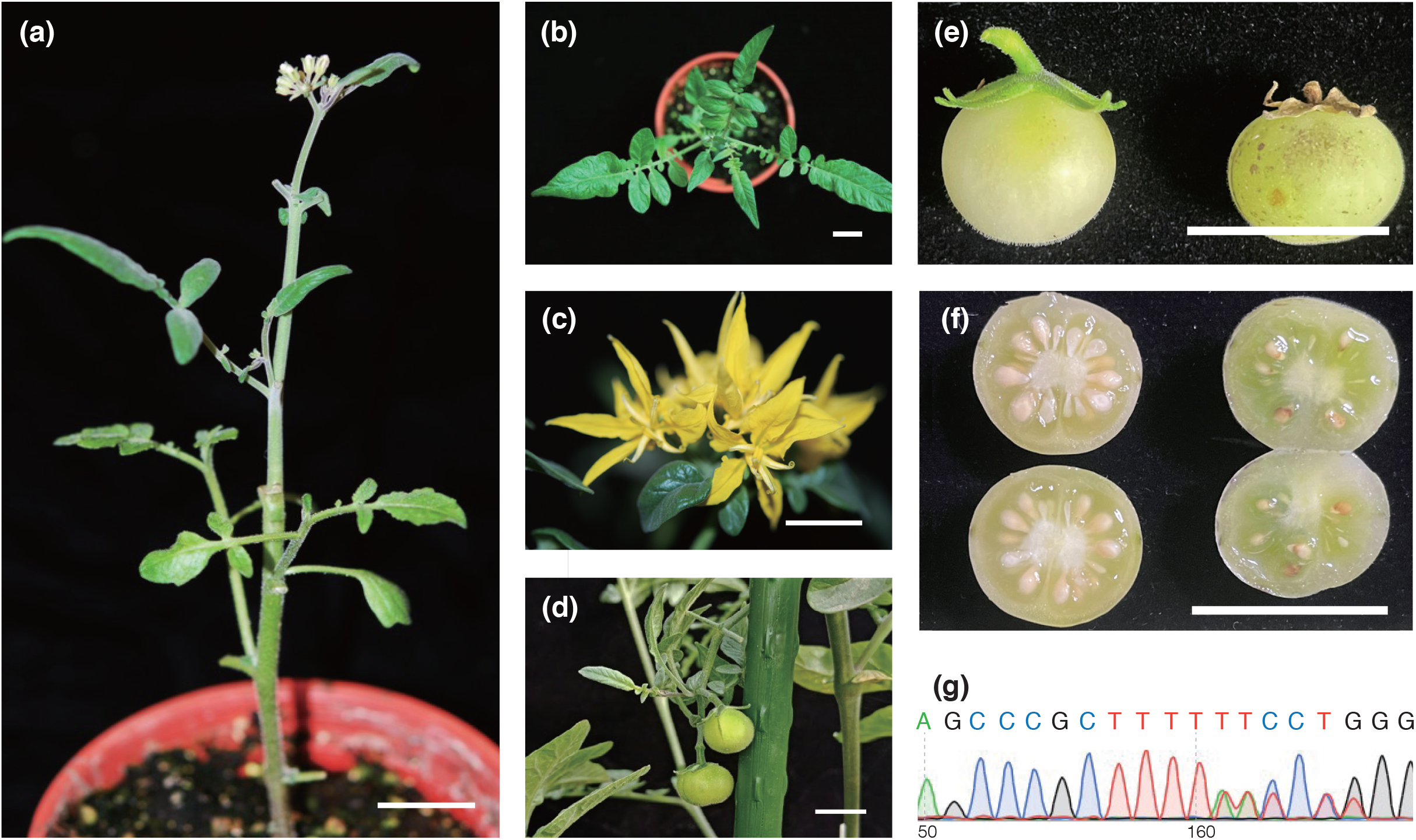
Growth of a sterile *spsgs3 #11* plant grafted with wild-type stock. **(a)** Grafted plant. Gray: null mutant. Leaves **(b)**, flowers **(c)**, and fruit of *spsgs3#11* scion. Mature fruit **(e)** and seeds **(f)** of wild-type stock (left) and *spsgs3#11* scion (right). **(g)** Results of Sanger sequencing of the seedling derived from *spsgs3#11* scion fruit, which is heterozygous, harboring *spsgs3#11* mutated alleles mixed with the wild-type allele. Bars = 1 cm.

## Discussion

In addition to its use in wild tomatoes, CRISPR is also utilized in commercial varieties of *S. lycopersicum*. Dozens of studies using this technique in tomato have been published, most involving breeding trials for traits such as quality (fruit architecture, color, metabolism, postharvest), anti-stress (biotic and abiotic stress), and domestication (Li et al., 2018; Zsogon et al., 2018). These studies were performed using several CRISPR platforms established in tomato, including (1) Cas9 (Brooks et al., 2014) and Cas12a (Bernabe-Orts et al., 2019), to generate DNA double-strand breaks that are preferentially repaired by non-homologous end joining to introduce target mutations; (2) precise modification of plant genomes using DNA repair templates via homologous recombination (Cermak et al., 2015); and (3) the cytidine base editor, an inactive Cas9 fusion with cytidine deaminase, which converts cytosine to uracil without cutting DNA and introducing mutations (Shimatani et al., 2017). Therefore, CRISPR is emerging as a powerful tool for tomato breeding.

For commercial breeding, however, it is desirable to produce DNA-free plants to avoid concerns about GMOs. Although there are several reports demonstrating successful DNA-free genome editing via biolistic methods in many crops (Svitashev et al., 2016; Liang et al., 2018; Banakar et al., 2020), protoplast regeneration systems have higher efficiency. Many studies have been performed using RNPs and plasmids to achieve DNA-free genome editing (Woo et al., 2015; Andersson et al., 2018; Lin et al., 2018; Hsu et al., 2019; De Bruyn et al., 2020; Hsu et al., 2021; Hsu et al., 2021; Yu et al., 2021). These reports indicate that it is possible to establish protoplast regeneration platforms for tomato and various target crops/plants.

Tomato and related species have been important materials in the development of protoplast isolation and regeneration techniques. Tomato protoplasts were isolated by enzymatic digestion, and this landmark achievement allowed sufficient amounts of protoplasts to be obtained for further application (Cocking, 1960). *S. peruvianum* was the first tomato-related species for which a protoplast regeneration system was reported, and such systems have subsequently been achieved in many tomato and wild tomato species (Kut and Evans, 1982). In this study, we combined these techniques to achieve DNA-free genome editing of a wild tomato. This method could be applied to other tomato-related species to facilitate breeding.

Although protoplast regeneration was first reported 50 years ago (Takebe et al., 1971), it still represents a major bottleneck in DNA-free genome editing. The major issue is that various species have different regeneration capacities. Moreover, no single protocol can be directly applied to all species efficiently because the requirements for plant medium and regeneration are diverse (Kut and Evans, 1982) and must be individually modified. Understanding how a cell is regenerated into a complete plant is an important topic of scientific and agricultural research (Maher et al., 2020), but information about this process is still limited. Such knowledge could be applied to develop efficient tissue culture, gene transformation, and genome editing system, tools that are important for *de novo* plant domestication (Li et al., 2018; Zsogon et al., 2018; Maher et al., 2020; Yu et al., 2021). In this study, we assessed the effects of plant growth regulators in the medium on protoplast regeneration. In addition to the chemical approach, several genes encoding morphogenic regulators have been identified and used to improve the efficiency of plant regeneration. It is possible to control the expression of these genes to establish a non-tissue-culture regeneration system for gene editing (Maher et al., 2020).

In addition to their roles in the domestication of wild species, polyploid crops have other benefits, including larger plants (Chung et al., 2017), and higher yields (Chen et al., 2018). In addition, triploid crop cultivars of species such as bananas and watermelons can produce commercially desirable seedless fruits. Most previous methods for chromosome multiplication have used colchicine. This procedure is complicated and inefficient, producing regenerated plants with mixed cell populations of various ploidy levels (Cola et al., 2014). Similar to haploid culture, in this report, using isolated protoplasts from polyploid cells in explants for regeneration and gene editing, we were able to obtain edited polyploid regenerated wild tomatoes without colchicine treatment. This phenomenon has also been reported in other plant species. In witloof chicory plants generated from CRISPR/Cas-edited protoplasts, 77.2% diploid and 21.5% tetraploid plants were produced and the remaining 1.3% consisted of haploids, hexaploids, and mixoploids (De Bruyn et al., 2020). Therefore, explants containing high proportions of polyploidized cells could be widely used for protoplast regeneration for crop polyploidization. However, in this study, we found no significant enlargement in the leaves or flowers of tetraploid versus diploid lines, similar to the pattern reported for tetraploid tomatoes (Nilsson, 1950).

In addition to technological difficulties, the presumed mutagenicity of protoplast regeneration is another reason why researchers are reluctant to use this system as a gene editing platform. Indeed, whole genome sequencing has revealed widespread genome instability in potatoes regenerated from protoplasts (Fossi et al., 2019), which has increased the concerns about this technology. The original purpose of protoplast regeneration was to use protoplast fusion to improve hybridization or as a platform for mutagenesis. Since only successful cases of mutation or fusion have been reported, and most such experiments have not been compared with other tissue culture methods, many researchers have the impression that protoplast regeneration readily leads to mutagenesis. In fact, other tissue culture technologies, including multiple shoot proliferation (Lin et al., 2007) and somatic embryogenesis (Lin et al., 2007), can also cause mutations. Although this study involved the use of PEG-Ca^2+^ in the transfection process, which could promote cell fusion, non-transfected tetraploid regenerated plants were also obtained. Based on our finding that the proportion of tetraploid regenerated plants was similar to that of shoot explants, we believe that the formation of polyploid regenerated plants was primarily due to the presence of polyploid cells in the explants. In addition to protoplast regeneration, there are also opportunities to obtain polyploid plants using other tissue culture technologies (Chung et al., 2017). In an *Agrobacterium*-mediated transformation experiment in tomato, the rate of tetraploid transgenic plants ranged from 24.5% to 80% and depended on both the genotype and the transformation procedure (Ellul et al., 2003). In *Arabidopsis* T-DNA insertion mutagenesis, large-scale genomic rearrangements have occurred (Pucker et al., 2021). Therefore, we believe that protoplast regeneration is an excellent tool for gene editing as well as other transgenic platforms.

Unlike the previous report of widespread genome changes in the autotetraploid potato (Fossi et al., 2019), the whole genome sequencing analysis in this study does not identify aneuploidy and abnormal chromosomal changes in either diploid or tetraploid regenerants. Chromosomes in the autotetraploid genome, such as cultivated potato, were derived from merging of two different chromosome sets (Van de Peer et al., 2021). On the other hand, tetraploid plants in this study, which were derived from chromosome doubling, contained the two identical sets of chromosomes. As the tissue culture steps caused a certain level of cell stresses, pairing of non-homologous chromosomes in the autotetraploid genomes (Fossi et al., 2019) likely has a higher probability of incorrect chromosome pairing than in the allotetraploid genomes (Hsu et al., 2019; Yu et al., 2021). Incorrect chromosome segregation during mitosis in the autotetraploid cells likely has a higher probability of evading the spindle-assembly checkpoint (Musacchio and Salmon, 2007). Furthermore, by analyzing changes in the allele frequency and copy number variation, we confirmed that the CRISPR-Cas9 editing did not introduce large scale chromosomal changes and unintended genome editing sites (Hsu et al., 2021).

In this study, all tetraploid and diploid *spsgs3* and *sprdr6* null mutants had wiry phenotypes, similar to other microRNA biogenesis null mutants in tomatoes (Yifhar et al., 2012; Brooks et al., 2014). *sgs3* and *rdr6* null mutants show various phenotypes in different species. *N. benthamiana spsgs3* and *sprdr6* mutants have a wiry flower morphology and sterile phenotype, but their leaves are similar to those of the WT (Hsu et al., 2021). The *Arabidopsis sgs3* mutant shows no significant phenotype (Adenot et al., 2006). Therefore, we would like to discover ways to improve the fertility of these mutants.

Grafting is a traditional agricultural tool that is used to control flowering, improve fruit quality, and increase resistance to biotic and abiotic stress (Haroldsen et al., 2012). In *N. benthamiana*, gene silencing was transmitted with 100% efficiency in a unidirectional manner from silenced stocks to non-silenced scions expressing the corresponding transgene (Palauqui et al., 1997). In this study, a mutant of *SpSGS3*, an RNA silencing-related gene, was used as a scion and grafted onto RNA-silenced normal wild-type rootstock. The fertility of *spsgs3#11* scions was rescued, and they produced seeds with mutated alleles. In *Arabidopsis*, more than 3,000 mobile genes have been identified. The mRNA from these genes could be transported long distance, including *SGS3* mRNA (Thieme et al., 2015). In addition to mRNA, organellar DNA, proteins, and plant growth regulators can also move across graft unions (Haroldsen et al., 2012). Whether these mobile substances were also involved in rescuing the fertility of the *spsgs3#11* scions or whether grafting with wild-type plants could rescue other sterile mutants of mobile RNA requires further investigation.

## Conclusions

To obtain tetraploid *S. peruvianum* DNA-free genome-edited plants, we used *in vitro-*grown shoots, which contain high proportions of tetraploid cells, as explants for protoplast isolation and regeneration. The medium components were optimized, and genome-edited regenerants were obtained within 6 months. This is the first study in *S. peruvianum* describing the use of both RNP and plasmid CRISPR reagents for DNA-free genome editing, yielding a targeted mutagenesis efficiency of 60% without the need for marker gene selection. Diploid and tetraploid heritable mutants were obtained for all pathogen-related genes targeted in this study, including *SpSGS3, SpRDR6, SpPR-1, SpProsys*, and *SpMlo1*, and the expected phenotypes were obtained. In comparative whole genome sequencing analysis, protoplast derived CRISPR-Cas9 edited plants, either diploid or tetraploid, showed stable genome structure. The proliferation of TYLCV, an important viral disease of tomato, was increased in *spsgs3* and *sprdr6* null mutants. The reproductive growth defect of the *SpSGS3* mutant was successfully rescued by grafting with WT stock. The protocols and materials described in this study will be useful for tomato breeding.

## Materials and methods

### Plant materials

Sterile *S. peruvianum* plantlets were propagated by cutting and growing them in half-strength Murashige and Skoog (1/2 MS) medium supplemented with 30 mg/L sucrose and 1% agar, pH 5.7. The plantlets were incubated in a 26°C culture room (12 h light/12 h in dark, light intensity of 75 µmol m−2 s−1). The plantlets were cut and subcultured in fresh medium monthly.

### Protoplast isolation and transfection

Protoplast isolation and transfection of *S. peruvianum* were performed following our previously published method with minor modifications (Hsu et al., 2019). Protoplasts were isolated from the stems and petioles of *in vitro-*grown plantlets. Five or more stems (approximately 5 cm/each, total 0.2-0.25 g) were used to isolate roughly 1 × 10^6^ protoplasts. These materials were place in a 6-cm glass Petri dish with 10 mL digestion solution [1/4 Murashige and Skoog (MS) liquid medium containing 1% cellulose and 0.5% macerozyme, 3% sucrose, and 0.4 M mannitol, pH 5.7] and cut into 0.5-cm-wide strips longitudinally. The material was incubated at room temperature in the dark overnight. The digested solution was diluted in 10 mL W5 (154 mM NaCl, 125 mM CaCl_2_, 5 mM KCl, 2 mM MES, and 5 mM glucose) solution and filtered through a 40-µm nylon mesh. The sample was centrifuged at low speed (360 × g) for 3 min to collect the protoplasts. The protoplasts were purified in 20% sucrose solution and washed three times with W5 solution. The protoplasts were transferred to transfection buffer (1/2 MS medium supplemented with 3% sucrose, 0.4 M mannitol, 1 mg/L NAA, and 0.3 mg/L kinetin, pH 5.7) and adjusted to a concentration of 3 × 10^5^ cells/mL.

The protoplasts were transfected with plasmids by PEG-mediated transfection (Woo et al., 2015; Lin et al., 2018). A 400-µL sample (1.2 × 10^5^ protoplasts) was combined with 40 µL of CRISPR reagent (DNA: 20-40 µg; RNP: 10 µg) and mixed carefully. The same volume (440 µL) of PEG solution was added to the sample, mixed, and incubated for 30 min. To end the reaction, 3 mL of W5 was added, and the sample was mixed well. Transfected protoplasts were collected by centrifugation at 360 × g for 3 min. The protoplasts were washed with 3 mL of W5 and centrifuged at 360 × g for 3 min. The target sites are shown in Table 1.

### CRISPR/Cas reagents

The SpCas9 vector for dicot transformation (pYLCRISPR/Cas9P35S-N) (Ma et al., 2015) was isolated using a Plasmid Midi-prep kit (Bio-Genesis). Preparation of Cas9 protein and sgRNA and Cas9 RNP nucleofection were performed according to Huang et al., 2020. Cas9 RNP complexes were assembled immediately before nucleofection by mixing equal volumes of 40 μM Cas9 protein and 88.3 μM sgRNA at a molar ratio of 1:2.2 and incubating at 37°C for 10 min.

### Protoplast regeneration

Pooled protoplast DNA was used as a template to amplify the target genes for validation by sequencing. The putatively edited protoplasts were transferred to 5-cm-diameter Petri dishes containing 3 mL 1/2 MS liquid medium supplemented with 3% sucrose, 0.4 M mannitol, 1 mg/L NAA, and 0.3 mg/L kinetin for plant regeneration. Calli formed from the protoplasts after 1 month of incubation in the dark. The calli were subcultured in a 9-cm-diameter Petri dish containing fresh medium with cytokinin for 3-4 weeks in the light. Calli that had turned green were transferred to solid medium containing the same plant growth regulators. The explants were subcultured every 4 weeks until shoots formed after several subcultures. The shoots were subcultured in solid rooting medium (HB1: 3 g/L Hyponex No. 1, 2 g/L tryptone, 20 g/L sucrose, 1 g/L activated charcoal, and 10 g/L Agar, pH 5.2) for adventitious roots formation.

### Analysis of the genotypes of regenerated plants

Two pairs of primers were designed to amplify the sgRNA-targeted DNA region for each target gene. The PCR conditions were 94°C for 5 min, 35 cycles of denaturing (94°C for 30 s), annealing (55°C for 30 s), and polymerization (72°C for 30 s), followed by an extension reaction at 72°C for 3 min. The PCR product was sequenced by Sanger sequencing to confirm mutagenesis. The multiple sequences derived from mutated regenerated plants were bioinformatically separated using Poly Peak Parser (http://yosttools.genetics.utah.edu/PolyPeakParser/; (Hill et al., 2014)) or further confirmed by sequential T/A cloning and sequencing. The primer sequences are listed in Table S7.

### Estimation of genome size

Fresh leaves were finely chopped with a new razor blade in 250 µL isolation buffer (200 mM Tris, 4 mM MgCl_2_-6H_2_O, and 0.5% Triton X-100) and mixed well (Dolezel et al., 2007). The mixture was filtered through a 40-μm nylon mesh, and the filtered suspensions were incubated with a DNA fluorochrome (50 μg/mL propidium iodide containing RNase A). The samples were analyzed using a MoFlo XDP Cell Sorter (Beckman Coulter Life Science) and an Attune NxT Flow Cytometer (Thermo Fisher Scientific). Chicken erythrocyte (BioSure) was used as an internal reference.

### Whole genome sequencing

Leaves of *S. peruvianum* regenerates were harvested and genomic DNA was extracted using two independent protocols. A nuclei isolation protocol (Sikorskaite et al., 2013) was used on the wild type (SpB) sample to recover higher quality and quantity of DNA samples. Briefly, nuclei were extracted by 36mM sodium bisulfite, 0.35M Sorbitol, 0.1M Tris-base, 5mM EDTA, 2M NaCl, 2% (w/v) CTAB, and 2 ml 5% N-lauroylsarcosine sodium salt. The genomic DNA was then extracted by chloroform-isoamyl alcohol (24:1), ethanol precipitation, and further cleaned up by DNeasy Blood & Tissue Kit (69504, Qiagen) and AMPure (Beckman Coulter). The other nine samples used the chloroform-isoamyl alcohol (24:1) for DNA extraction, followed with Zymo Genomic DNA Clean & Concentrator-25 (D4064, Zymo), and Zymo OneStep PCR Inhibitor Removal Kit (D6030, Zymo) to obtain high quality genomic DNA. DNA integrity was checked using the D1000 Screen Tape on the Agilent TapeStation 4150 System with DIN value > 8. Genomic DNA were sheared using a Covaris E220 sonicator (Covaris) and paired-end sequencing libraries were constructed by the NEBNext Ultra DNA Library Prep Kit II for Illumina (E7370S, NEB). DNA libraries were validated again on the Agilent TapeStation 4150, and were quantified by qPCR (E7630, NEB). The 2×150 bp paired-end sequencing with average insert size of 700 bp was performed by Welgene Biotech on an Illumina NovaSeq 6000 platform.

### WGS data analysis

Since there was no assembled *S. peruvianum* genome, high quality Illumina reads were mapped to the *S. lycopersicum* Heinz 1706 reference genome (SL4.0) (Hosmani et al., 2019) by the GPU-based NVIDIA Clara Parabricks package (NVIDIA). To determine the variant frequency, we used the deep learning-based Google DeepVariant (Yun et al., 2021) with ‘WGS model’ to identify variants. All samples were then combined by GLnexus (Yun et al., 2021) to perform ‘joint genotype calling’ using ‘DeepVariant’ model to combine samples. We then calculated the heterozygous allele frequency by dividing the read depth of the heterozygous allele (labeled as 0/1 by GLnexus) over the total read depth of the variant. A large chromosomal region with heterozygous allele frequency lower than 0.5 indicated either the chromosome region with low recombination rate or deletion of the chromosome fragments. To determine CNVs between samples, we used the cn.mops pipeline (Klambauer et al., 2012) to analyze mapped Illumina reads. To minimize the effects of repetitive sequence regions, we set the segment size to 3,000 bp and minimum number of segments as 10 to identify high confidence CNVs.

### Quantitative real-time PCR (RT-qPCR)

Expression of four genes was analysed using real-time PCR. These genes were: *SpSGS3, SpARF3, SpARF4*, and *SpRDR6*. Transcripts of all four genes were profiled with three biological replications and each with at least three technical replications using the RNA samples of regenerants. RT-qPCR was carried out in 96-well optical reaction plates using the iQ™ SYBR^®^ Green Supermix (Bio-Rad). The reference gene Actin and gene-specific primers for the RT-qPCR are listed in Supplementary Table S8.

## Acknowledgments

We thank Te-Chang Hsu and the AS-BCST Bioinformatics Core for the computational support and Miranda Loney and Plant Editors for English editing. Experiments and data analysis were performed in part using the confocal microscope at the Division of Instrument Service of Academia Sinica with the assistance of Shu-Chen Shen. We thank IPMB Flow Cytometry Analysis and Sorting Service of Academia Sinica for flow cytometry analysis. We thank Ruei-Shiuan Wang for genomic DNA preparation, Yu-Jung Cheng for tissue culture, Wei-Fong Hung for genotyping, Ting-Li Wu and Jheng-Yang Ou for figure preparation and Song-Bin Chang for karyotyping.

## Author contributions

C-SL, Y-CL, JS, and M-CS conceived and designed the experiments. C-TH, and Y-HY performed the CRISPR-Cas9 experiments. C-TH, Y-HY, Q-WC, J-JY, and F-HW conducted the protoplast regeneration, cell biology, molecular biology, and targeted mutagenesis experiments. SL conducted SpCas9 purification. Y-LW performed WGS library preparation and qPCR analysis. P-XZ and Y-CL performed bioinformatics analysis. Y-HC, C-TH, C-SL, Q-WC, and F-HW performed virus-related analysis. C-TH performed cell biology. C-TH and S-IL performed grafting. JS, M-CS, Y-CL, and C-SL wrote the manuscript with input from all co-authors. All authors read and approved the final manuscript.

## Data Availability Statement

The Illumina sequencing reads generated for this study have been deposited at NCBI under BioProject PRJNA768623. (https://dataview.ncbi.nlm.nih.gov/object/PRJNA768623?reviewer=m0ufmjvjdqsj9evnf4ek6qtl8l)

## Supplemental Figures

**Figure S1. Effects of cytokinins on callus induction (1**^**st**^ **subculture) and callus proliferation (2**^**nd**^ **subculture)**. The effects of cytokinins [kinetin, zeatin, 6-(γ,γ-Dimethylallylamino)purine (2ip), and 6-Benzylaminopurine (BA)] during these two stages were investigated separately. Different cytokinins were added during callus induction [1^st^ subculture, 1/2 Murashige and Skoog (MS) medium supplemented with 3% sucrose, 0.4 M mannitol, pH 5.7 liquid medium supplemented with 0.2 mg/L cytokinin and 1 mg/L NAA]. Kinetin yielded the fewest calli, and the three other cytokinins led to better callus induction. During callus proliferation [2^nd^ subculture, 1/2 MS medium supplemented with 3% sucrose, 0.4 M mannitol, pH 5.7 liquid medium supplemented with 2 mg/L cytokinin and 0.3 mg/L Indole-3-acetic acid (IAA)], the addition of zeatin, 2ip, and BA caused the callus to grow and turn green. Inclusion of 2ip during callus induction yielded the same number of cells as the other cytokinin treatments, but the cell clusters were smaller and did not grow easily when directly transferred to callus proliferation medium in the light. Therefore, zeatin and BA are the best treatments for liquid culture. Bar = 1 cm.

**Figure S2. Effects of cytokinins on callus in solid medium (3**^**rd**^ **subculture)**. Calli from media containing different cytokinins (2^nd^ subculture) were transferred to solid medium containing the same cytokinin (3^rd^ subculture). Cytokinin in the medium had a strong effect on callus growth (Figure S4). Regardless of the callus induction medium used, browning of the callus occurred in solid medium supplemented with kinetin. Callus derived from 2ip callus induction medium proliferated only in 2ip solid medium. BA and zeatin had similar effects on callus growth, but calli on zeatin medium showed more greening. We therefore identified zeatin as the most suitable cytokinin for use in solid medium. Bar = 1 cm.

**Figure S3. Flow cytometric analysis of the nuclear DNA contents of tetraploid plants regenerated from *SpProsys* RNP-transfected protoplasts**. The number of regenerated plants is shown at the top left of each panel. Gray font: null mutant. The genome sizes are shown at the top right. The results are derived from three technical repeats. Unit: pg. Un-edited: The *SpProsys* sequences are similar to the wild type. Chicken erythrocyte nuclei (CEN: 2.5 Gb) were used as the calibration standard. The bar indicates the area used to count nuclei. The genome sizes of all seven regenerants were measured by flow cytometry, including two un-edited, three heterozygous, and two biallelic plants that were tetraploid. Both tetraploid and diploid regenerants (Table S7) derived from *SpProSys* RNP transfections flowered normally, and no distinctive phenotype was observed. Bar = 1 cm.

**Figure S4. Phenotypes of diploid and tetraploid plants regenerated from protoplasts transfected with CRISPR reagents**. Underline: 4*n*. Bars = 1 cm. *SpSGS3*#10, *SpSGS3*#7 and *SpRDR6*#38 contained mutated alleles. (a) the fruits of diploid and tetraploids regenerated from transfected protoplasts. (b) T_1_ seeds of the heterozygous diploid (*SpSGS3*#10) and tetraploid (*SpSGS3*#7 and *SpRDR6*#38) mutants. (c) 1.5-month-old T_1_ seedling derived from T_0_ transfected protoplast regenerated plants.

**Figure S5. Progeny analysis of *SpSGS3***. Underlined regenerated plant name: tetraploid. Red font: mutated nucleotide. Green/blue font: sequences shown in the green/blue boxes in the Sanger sequencing results. WT: wild type. M: mutant. WT:M: wild type/mutant ratio based on Sanger sequencing results. No.: number of progeny in this ratio. (a) *SpSGS3*#7 T_1_ progeny analysis. The allele sequences in the GTTCCTCCTGCTCTGAAGAA target site are listed; 0–3 mutated alleles were identified. This regenerated plant was shown to be allotetraploid. (b) The PCR product of the *spsgs3#7-2* null mutant was subjected to T/A cloning, and the clones were subjected to Sanger sequencing (GCGCAATTGAATGGTTTACA target site and ATTCCCCCCAGGATAAAAGC target site). Three types of mutated alleles were identified. (c) Analysis of diploid *SpSGS3*#10 T_1_ progeny.

**Figure S6. Progeny analysis of *SpRDR6***. Underlined regenerated plant name: tetraploid. Red font: mutated nucleotides. Blue font: sequences shown in blue boxes in the Sanger sequencing results. (a) *SpRDR6*#6-2 genotyping. Top: allele sequences. Middle: The Sanger sequencing results indicate the presence of multiple peaks after TTAAGCT. Bottom: The T/A cloning results demonstrate that *SpRDR6*#6-2 contains a mutated allele (M) similar to *SpRDR6*#6. (b) RT-PCR product of the *sprdr6*#33-G null mutant. The result indicates that *sprdr6*#33-G is a homozygous null mutant. The mutated allele can still generate a transcript. (c) Genotyping of the *sprdr6*#38-6 null mutant. Top: The allele sequences of *SpRDR6*#38. Middle: Sanger sequencing results of *sprdr6*#38-6 genomic DNA. Bottom: The M1 and M2 mutated alleles identified by T/A cloning without wild-type alleles.

**Figure S7. Progeny analysis of *SpPR-1***. Red font: mutated nucleotide(s). Blue/green font: sequences shown in blue/green boxes in the Sanger sequencing results. (a) Progeny analysis of *sppr-1*#52. Top: allele sequences. Middle: Sanger sequencing results of different genotypes. Multiple peaks are shown in heterozygous lines (M1M2, M1M3, M2M3). No.: number of progeny of each genotype. Bottom: M3 sequence identified by T/A cloning. (b) Progeny analysis of *sppr-1*#61. Top: allele sequences. Middle: *SpPR-1* genomic PCR products of *sppr-1*#61 progeny. The genotypes of individual progeny were determined based on DNA size and are shown below the image. Sanger sequencing results for the LL and SS genotypes.

**Figure S8. Illumina sequencing coverage for the tomato SL4.0 genome assembly**. The Illumina PE reads were mapped by BWA and the sequencing depth was calculated in 10kb window size. Coverage is plotted using 30 bins per chromosome on the X axis. Black dashed line: median of the sequencing coverage of each chromosome.

**Figure S9. Phenotypes of the *spsgs3* and *sprdr6* null mutants**. Underlined regenerated plant name: tetraploid. Wiry phenotypes of T_0_ diploid *spsgs3* null mutants #6 and #13. Bar = 1 cm. (b) Wiry phenotypes of T_1_ tetraploid *spsgs3*#7-2 and *sprdr6*#38-16. Bar = 1 cm. (c) Alexander staining of wild-type and *spsgs3*#11 pollen. Bar = 50 μm.

**Figure S10. Symptoms and TYLCV proliferation on *in vitro*-cultured *S. peruvianum* plants inoculated with the infectious TYLCV clone**. Gray: null mutant. Underline: 4*n*. Bars = 1 cm. *SpRDR6*#2 and *SpSGS3*#24 were non-mutated protoplast regenerated plants. Line 1, 7: *SpRDR6*#2; 2, 8: *sprdr6*#38-6; 3, 9: Wild type; 4, 10: *spsgs3*#11; 5, 11: *SpSGS3*#24; 6, 12: *spsgs3*#7-2

## Supplemental Tables

**Table S1. *SpSGS3* gene sequences of the *SpSGS3* mutants**. Gray: null mutant. Underline: 4*n*. Red font: mutated nucleotide(s). Number in brackets: length of nucleotide sequence. −: deletion. +: insertion.

**Table S2. *SpRDR6* gene sequences of the *SpRDR6* mutants**. Underline: 4*n*. Red: mutated nucleotide(s). −: deletion. +: insertion.

**Table S3. *SpPR-1* gene sequences of the *SpPR-1* mutants**. Gray: null mutant. Underline: 4*n*. Red: mutated nucleotide(s). Number in brackets: length of nucleotide sequence. −: deletion. +: insertion.

**Table S4. *SpProsys* gene sequences of the *SpProsys* mutants**. Gray: null mutant. Underline: 4*n*. Red: mutated nucleotide(s). Number in brackets: length of nucleotide sequence. −: deletion. +: insertion.

**Table S5. *SpMlo1* gene sequences of the *SpMlo1* mutants**. Gray: null mutant. Underline: 4*n*. Red: mutated nucleotide(s). Number in brackets: length of nucleotide sequence. −: deletion. +: insertion.

**Table S6. Karyotypes of plants regenerated from protoplasts transfected with CRISPR reagents**. WT: the target gene sequences are un-edited, like the wild type. *: the genome size was determined by flow cytometry.

**Table S7. Overview of Illumina WGS sequencing, mapping rate and SRA number**.

**Table S8. Primers used in these studies**.

